# Discovering complete quasispecies in bacterial genomes

**DOI:** 10.1101/119743

**Authors:** Frederic Bertels, Chaitanya S. Gokhale, Arne Traulsen

## Abstract

Mobile genetic elements can be found in almost all genomes. Possibly the most common non-autonomous mobile genetic elements in bacteria are REPINs that can occur hundreds of times within a genome. The sum of all REPINs within a genome are an evolving populations because they replicate and mutate. We know the exact composition of this population and the sequence of each member of a REPIN population, in contrast to most other biological populations. Here, we model the evolution of REPINs as quasispecies. We fit our quasispecies model to ten different REPIN populations from ten different bacterial strains and estimate duplication rates. We find that our estimated duplication rates range from about 5 × 10^−9^ to 37 × 10^−9^ duplications per generation per genome. The small range and the low level of the REPIN duplication rates suggest a universal trade-off between the survival of the REPIN population and the reduction of the mutational load for the host genome. The REPIN populations we investigated also possess features typical of other natural populations. One population shows hallmarks of a population that is going extinct, another population seems to be growing in size and we also see an example of competition between two REPIN populations.

## Introduction

Repetitive sequences are common in most bacterial genomes, but rare compared to most eukaryotic genomes (Jurka *et al*. 2007; Versalovic *et al*. 1991). A large proportion of repetitive sequences in bacterial genomes are the result of self-replicating DNA sequences. These sequences usually encode an enzyme called a transposase that specifically copies its own sequence (Mahillon and Chandler 1998). There are also repetitive sequences that do not encode a transposase themselves, but are copied by a transposase that is encoded elsewhere in the genome. These elements are referred to as MITEs (<u>M</u>iniature <u>I</u>nverted repeat <u>T</u>ransposable <u>E</u>lements) (Wessler *et al*. 1995). MITEs were first described in plant genomes (Bureau and Wessler 1994) and later also in bacteria (Oggioni and Claverys 1999). Recently, it has been shown that REP (<u>R</u>epetitive <u>E</u>xtragenic <u>P</u>alindromic) sequences (Higgins *et al*. 1982) or more specifically REPINs (<u>R</u>EP doublets forming hairp<u>IN</u>s) (Bertels and Rainey 2011b), one of the most abundant repeat families in bacteria, are also MITEs (Nunvar *et al*. 2010; Bertels and Rainey 2011b, a; Ton-Hoang *et al*. 2012).

REP sequences are about 25 bp long sequences that are highly abundant in bacterial genomes (Higgins *et al*. 1982; Aranda-Olmedo *et al*. 2002; Silby *et al*. 2009). They contain a short imperfect palindromic sequence that can form short hairpins in single stranded DNA or RNA. REP sequences mostly occur in non-coding DNA between genes and are part of REPINs. REPINs in most *Pseudomonas* strains consist of two REP sequences in inverted orientation separated by a highly diverse nucleotide sequence (Bertels and Rainey 2011b). REPINs are a replicative unit and are mobilized by RAYTs (<u>R</u>EP <u>A</u>ssociated t<u>Y</u>rosine <u>T</u>ransposases) (Nunvar *et al*. 2010; Bertels and Rainey 2011b; Ton-Hoang *et al*. 2012). Although the structure of REPINs in *Pseudomonas* is well defined, for REPINs in *E. coli* there has not been an extensive study on what exactly comprises the replicative unit.

The occurrence of REP sequences and associated functions have been described in many different bacterial genomes (Higgins *et al*. 1982; Aranda-Olmedo *et al*. 2002; Silby *et al*. 2009). However, their evolution has rarely been studied in detail (Bertels and Rainey 2011a, b) and nothing is known about the duplication rates of REPINs. Although, we know that closely related E. *coli* strains contain varying numbers of REP sequences, this may not be a direct result of replication. Instead it may be more likely that it is a consequence of the extremely dynamic genome composition of E. *coli* (Touchon *et al*. 2009), where REP sequences get deleted or inserted together with other parts of the genome. However, the lack of evidence for novel REPIN insertions probably means that duplication rates are low, despite the presence of hundreds of REPINs in some genomes (Bertels and Rainey 2011b).

As it is difficult to study the evolution of the complete REPIN sequence due to the highly diverse loop region (which is probably strongly affected by recombination), we model the evolution of the most conserved 25bp at each end of the REPIN. Here we infer REPIN duplication rates by modeling the most abundant REPINs in a bacterial genome as a quasispecies in equilibrium. The beauty of studying REPINs in bacterial genomes is that we know the exact composition of the population at the time of genome sequencing, something that is impossible to achieve for almost any other population study.

We first fit the equilibrium of our quasispecies model for a REPIN population from *Pseudomonas fluorescens* SBW25 and later for nine other bacterial genomes. Our results show that despite the large divergence between the bacterial strains, our inferred duplication rates are very similar and very low. All rates fall into a narrow margin between one replication in about 31 × 10^6^ and 200 × 10^6^ host divisions. Hence, if a bacterium were to divide every 40 minutes, it would take about 2359 years for a specific REPIN duplication to fix in the population. The astonishing rarity of these events may explain the lack of evidence for novel REPIN insertions in bacterial genomes.

## Materials and Methods

### Quasispecies model

The quasispecies model describes the mutation-selection balance of a set of similar sequences that evolve on a fitness landscape. On this landscape, each sequence has a certain fitness. Sequences with high fitness leave many offspring, sequences with low fitness leave few offspring. The fitness landscape is traversed by acquiring mutations (Eigen 1971; Eigen and Schuster 1977; Nowak 1992).

The quasispecies model has been applied previously mostly to model viral populations (Seifert *et al*. 2015; Domingo and Schuster 2016). Here, we model REPIN sequences that mutate and duplicate: the fitness in the quasispecies model corresponds to the REPIN duplication rate and the model’s mutation rate to the genome mutation rate. We assume that the REPIN population in our genome is a quasispecies in equilibrium. The most abundant sequence in our population is our master sequence. With increasing genetic distance to the master sequence, fitness changes. For our model we assume five discrete fitness classes. The 0^th^ class contains the master sequence. Sequences differing in 1, 2 or 3 positions are in the next three classes. The remaining sequences are in the 4^th^ fitness class. The frequencies of the sequences belonging to each of these classes *i* are given by *x_i_*. The population evolves to a mutation-selection balance as described by the standard quasispecies equation (Page and Nowak 2002; Bull *et al*. 2005)

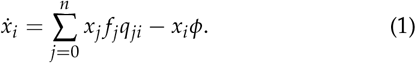

In our case *n* equals 4. The fitness of sequences belonging to each class *j* is given by *f_j_* and the average fitness of the population by 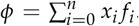. The probability that a sequence from class *j* mutates into *i* is given by *q_ji_*. In our model, sequences can only acquire a single mutation per time step. Hence, **Q** is a tri-diagonal matrix with non-zero entries in the main diagonal (no mutation) the first diagonal above (sequence acquires an additional mutation) and the first diagonal below (back mutation). For a mutation rate *μ* and a sequence length *L*, the probability of transitioning to the next mutation class *i* + 1 is 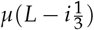 and to the previous mutation class *i* – 1 is 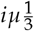. The fourth mutation class is the only class where we assume a back mutation rate of zero — the exact value would depend on the frequency distribution of the sequences that differ by more than three mutations to the master sequence. We also assume that the mutation rate of REPINs only depends on the host mutation rate. Mutations that occur during the duplication process are assumed to be negligible.

### Parameterizing the quasispecies model

We set the fitness of the highest mutation class to one, *f*_4_ = 1. For a given set of equilibrium sequence frequencies, we can then calculate the relative fitness of the remaining four mutation classes for a given mutation rate (see File S7). For all our bacteria we assume a host mutation rate of 8.9 × 10^−11^, which was inferred for *E. coli* (Wielgoss *et al*. 2011). The duplication rate is then the calculated fitness for each mutation class subtracted by one.

### Stochastic simulations

For each REPIN population, we performed a stochastic simulation to determine the extent of stochastic fluctuation on the equilibrium frequencies. These fluctuations mainly depend on the REPIN population size. As we cannot simulate evolution for the genome mutation rate, we scaled our fitness values up to fit a mutation rate of 10^−4^. With the new mutation rate, each discrete time step corresponds to 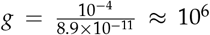 bacterial generations. Because we assume multiplicative fitness, the fitness values at a mutation rate of 10^−4^ are comparable to (*f_i_*)^*g*^.

We modeled evolution with a Wright-Fisher process (Ewens 1979). We start the simulation with a clonal population of the master sequence at carrying capacity, which is set to the number of REPINs observed in the genome. The number of offspring each sequence leaves in each generation is equal to the sequence’s fitness. If the number of offspring exceeds the carrying capacity, a random selection of the same size as the carrying capacity survives to the next time point. We modeled a total of 10^5^ generations.

We repeated each simulation 100 times and measured the proportion of simulations where the 0^th^ mutation class persisted at a frequency of more than 10%.

### Determining REPIN populations

We extracted REPIN populations from 10 bacterial genomes the following way: For each of these genomes we determined the most common 25 bp long sequence. We then recursively searched the genome for all sequences that have a Hamming distance of 2 to all identified sequences until no more sequences were found. We call these sequences REP sequences. For all REP sequences we determined whether they were part of a sequence cluster by checking whether there were any additional occurrences in a vicinity of 130bp. From these sequence clusters we extracted REPINs. REPINs consist of two adjacent REP sequences that are found in opposite directions (one on the positive strand the other on the negative DNA strand, also called inverted repeats) in the DNA sequence. The REPINs we found were extracted and joined together facing the same direction in alphabetical order. REP sequences found as direct repeats or as singlets in the genome were also extracted (as single sequences). We added another 25bp of adenine nucleotides at the end of each REP singlet to make them easily comparable with REPINs.

### Clustering REPIN sequences

REPIN populations can be represented as sequence networks. In these networks, each node represents a sequence. Vertices between nodes exist if the Hamming difference between the sequence pair is one. Because REPIN populations in *Pseudomonas* do not always evolve on a single peak due to the presence of multiple RAYTs (transposases) in the genome, we extracted subpopulations clustered around the master sequence. We determined these subpopulations for all *Pseudomonas* strains by applying a Markov clustering algorithm implemented in the MCL package (van Dongen 2000) with the inflation parameter set to 1.2 to the sequence network. The MCL algorithm simulates random walks on a stochastic graph by alternating between expansion and inflation operations, where larger inflation parameters will lead to more fragmented networks

We used the largest REPIN cluster for our analyses. Since these clusters exclude decayed sequences far from the master sequences, we also included all sequences with a Hamming distance of two to any sequence in the cluster. Of the sequences identified in the last step we only included instances that occurred less than three times in the genome. Sequences that occur more than three times in the genome are likely to have been duplicated by other RAYTs.

### Inferring an error threshold

The error threshold defines a critical point in a quasispecies where with the given fitness values and mutation rate it is impossible to maintain the master sequence. Here we deviate slightly from this definition as we define the error threshold as the point where the master sequence cannot be maintained at a relative frequency of more than 1%. To determine the duplication rate at which we reach our error threshold, we decrease all fitness values in increments of 1 × 10^−12^. As soon as one of the five fitness parameter reaches one, this parameter will remain constant for the remainder of the procedure. We performed this procedure for the fitness landscape of each species separately.

### Data Availability

All genomes are publicly available on Genbank (https://www.ncbi.nlm.nih.gov/genbank/) under the following accession numbers:

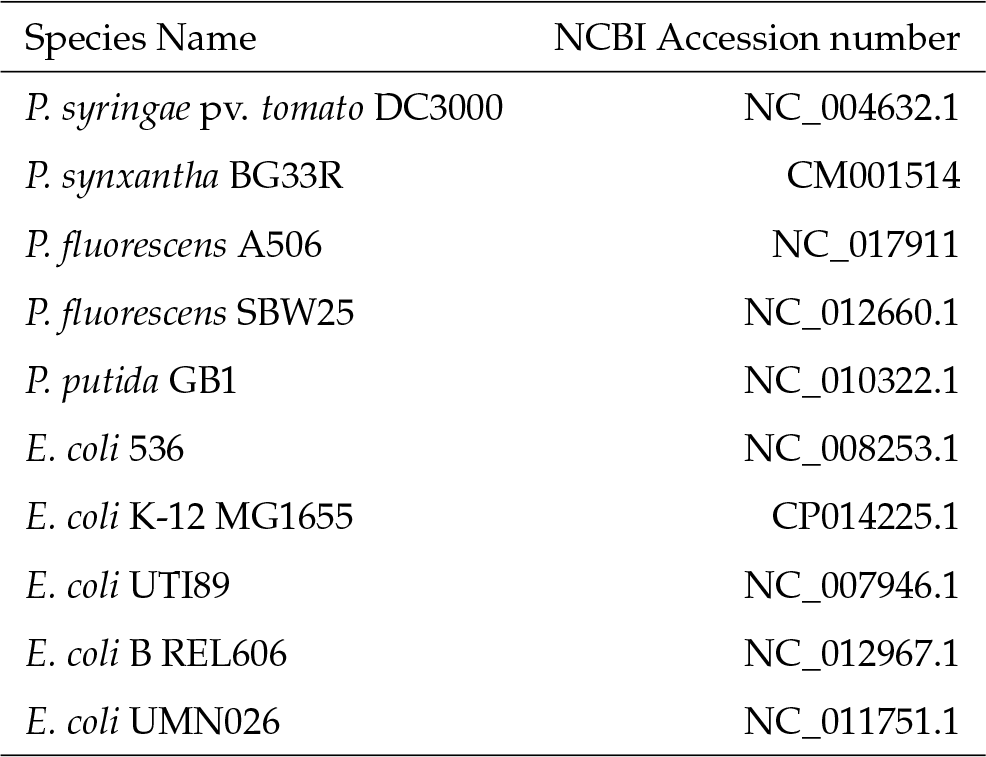

We included eight supplemental files. File S1 contains detailed descriptions of all supplemental files. File S2 contains the sequence and frequency of the most common 25 bp long sequence, the gene name of the flanking RAYT and the number of RAYTs, in all of the bacteria analyzed in this study. File S3 contains the modeling and simulation results for all ten REPIN populations we analyzed in our study. File S4 contains the Proportion of symmetric REPINs in all identified sequences from all studied strains. File S5 contains the duplication rates and equilibrium frequencies for each of the 10 REPIN populations at the error threshold. File S6 contains the Mathematica code we used to calculate equilibrium frequencies, fitness values and error thresholds for all 10 REPIN populations. File S7 contains the same Mathematica code as pdf. File S8 contains the sequence frequencies of the different mutation classes for all 10 REPIN populations.

## Results and Discussion

### *REPINs in* Pseudomonas fluorescens *SBW25*

In *Pseudomonas fluorescens* SBW25 REPINs consist of two inverted highly conserved sequences that are 25 bp in length, separated by a sequence of varying length that shows low levels of conservation (Bertels and Rainey 2011b, a). The processes that lead to the varying levels of conservation in REPINs are not very well understood. Hence, we will focus our analysis only on the most conserved 25 bp flanking the REPIN. These sequences have been discovered a long time ago in *E. coli* and have been called REP sequences (Stern *et al*. 1984). To find the most conserved parts of the REPIN, we determined the most common 25bp long sequence in the SBW25 genome. This sequence occurs 265 times and is usually part of a REPIN (Bertels and Rainey 2011b). We then add all sequences that differ in no more than two positions to this sequence. For the identified sequences we do the same and so on, until we can find no more new sequences in the genome. The resulting REP population contains 932 REP sequences. For these sequences, we determine whether they are part of a REP cluster, by looking for all occurrences in the vicinity of 130bp. From these clusters, we extract adjacent pairs of inverted REP sequences or REPINs. REP singlets were also extracted but marked with a 25bp long adenine sequence. The relationship between REPINs is visualized as a sequence network (Figure 2).

**Figure 1.**
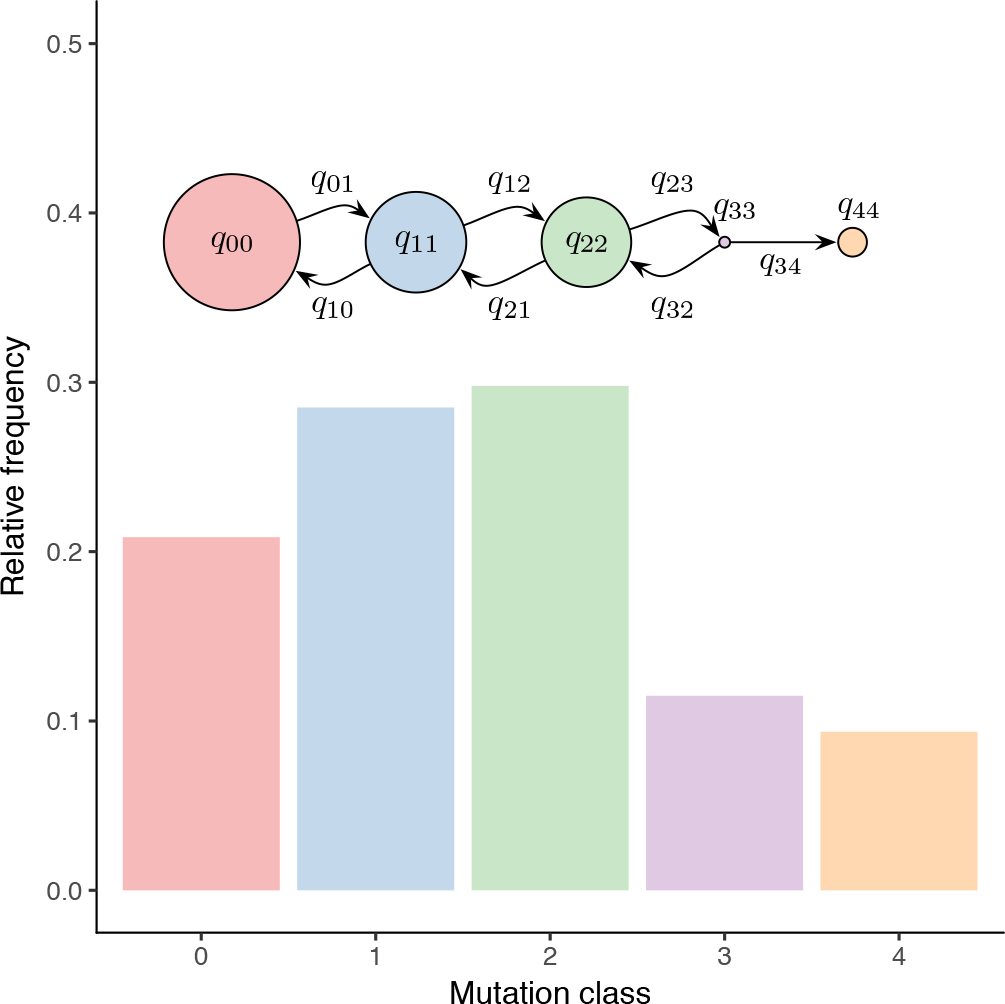
Exemplar results for a quasispecies model. For a mutation rate of *μ* = 8.9 × 10^−11^, and the fitnesses as given in Table 1 (1+scaled duplication rate), we illustrate the equilibrium distribution of the relative frequencies of *P. fluorescens* SBW25 REPINs. The radii of the circles indicate the duplication rate, which is the quasispecies fitness subtracted by one. Note that the actual fitness differences are extremely minute at the level of 10^−9^. The cartoon merely illustrates the architecture of the fitness landscape. The mutation probabilities are given by (*q_ij_*) while self-replication occurs with probability *q_ii_*.

**Figure 2.**
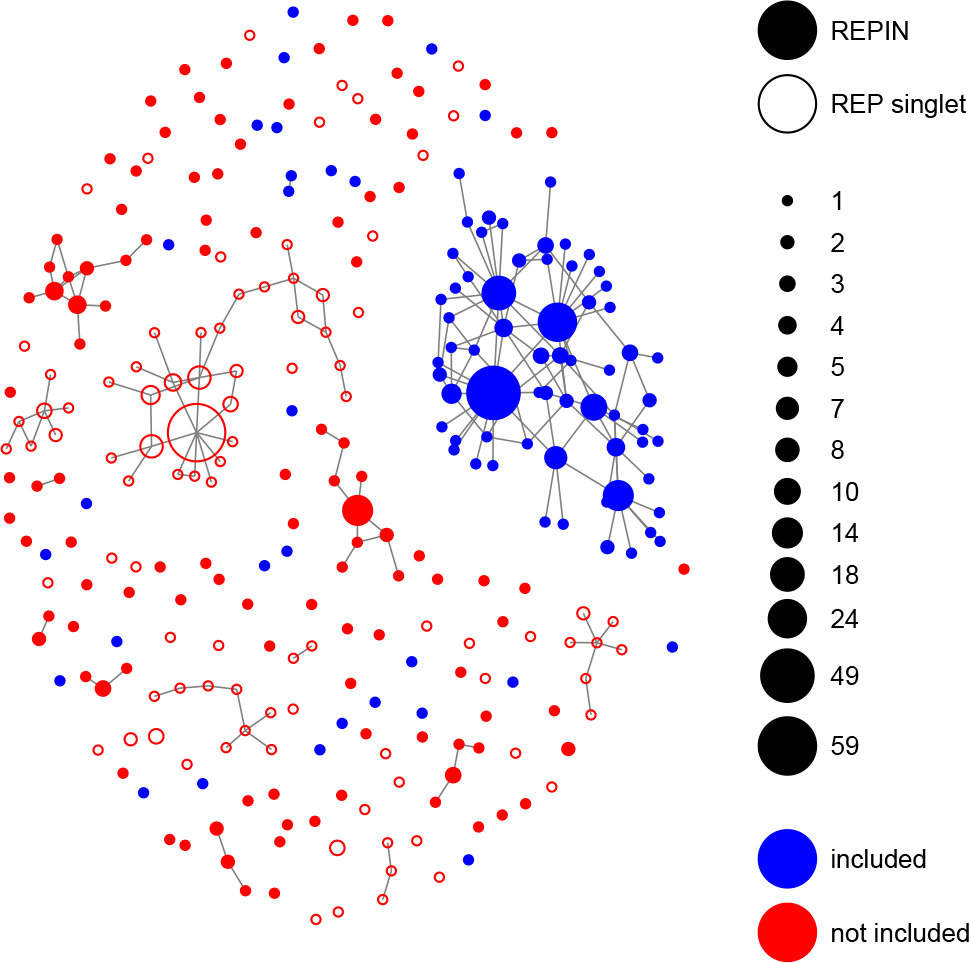
Structure of the REPIN population in SBW25. REPINs that differ in exactly one position are connected. REP sequences that do not form REPINs (e.g. singlets) are shown as empty circles. Blue “included” nodes belong to the REPIN population for which we infer duplication rates. Red (“not included”) nodes were excluded from the analysis because they likely evolve on a more complex fitness landscape that is more difficult to model. The size of the nodes indicates the frequency of the corresponding sequence in the SBW25 genome.

The population network in Figure 2 has many sequence hubs distantly related and not connected to the master sequence. Instead of a very rugged activity landscape of a single RAYT (the transposase responsible for duplicating REPINs), we think it is more likely that these hubs were created by the concurrent activities of multiple RAYT transposases (the SBW25 genome contains three RAYT genes). As it is impossible to accurately model this complexity for small REPIN populations, we decided to reduce the REPIN population to all sequences that are part of the largest cluster as well as all sequences that are at most 2bp different from any sequence that is part of the cluster.

The “included” subpopulation selected in Figure 2 has 235 members. We will model this subpopulation as quasispecies, with five sequence classes, that are 0, 1, 2, 3 and more than 3 mutations away from the master sequence. In our model we will also assume that the population is in equilibrium and the frequencies of the sequences we observe are steady state frequencies. The mutation rate in our model was chosen to be high to facilitate stochastic simulations of the evolutionary process. The fitness values for each mutation class were calculated from the quasispecies equation for the sequence frequencies observed in SBW25 (Table 1).

**Table 1.** Inferred REPIN duplication rates in *P. fluorescens* SBW25.

| Mutation class | Inferred Duplication Rate $\lambda_i (\times 10^{-9})^a$ | Scaled Duplication Rate $\tilde{\lambda}_i (\times 10^{-9})^b$ |
| --- | --- | --- |
| 0 | 9.8 | 11.3 |
| 1 | 6.5 | 8.1 |
| 2 | 5.5 | 7.1 |
| 3 | -1.6 | 0 |
| 4 | 0 | 1.6 |

<sup>a</sup> We identified a master sequence in the data and inferred the frequency of the different mutation classes. We use the equilibrium of our quasispecies model to calculate the associated fitness values $f_i$ and setting $f_4$ to 1, where $\lambda_i$ is $f_i - 1$ .
<sup>b</sup> The scaled duplication rate is: $\tilde{\lambda}_i = \frac{f_i}{\min(f_i)} - 1$ .

The quasispecies equation provides us with a set of fitness values that perfectly recapitulate the observed frequencies for infinitely large populations (Figure 3A). However, REPIN populations are relatively small, which means that population size will have a strong effect during REPIN evolution. To estimate stochastic effects, we used the calculated fitness parameters for each mutation class to perform a stochastic Wright-Fisher simulation with a maximum of 235 individuals (Figure 3B). Our simulation shows that the distributions of the mutation classes are wide, particularly for the master sequence, which is probably an effect of the small population size (Figure 3C).

**Figure 3.**
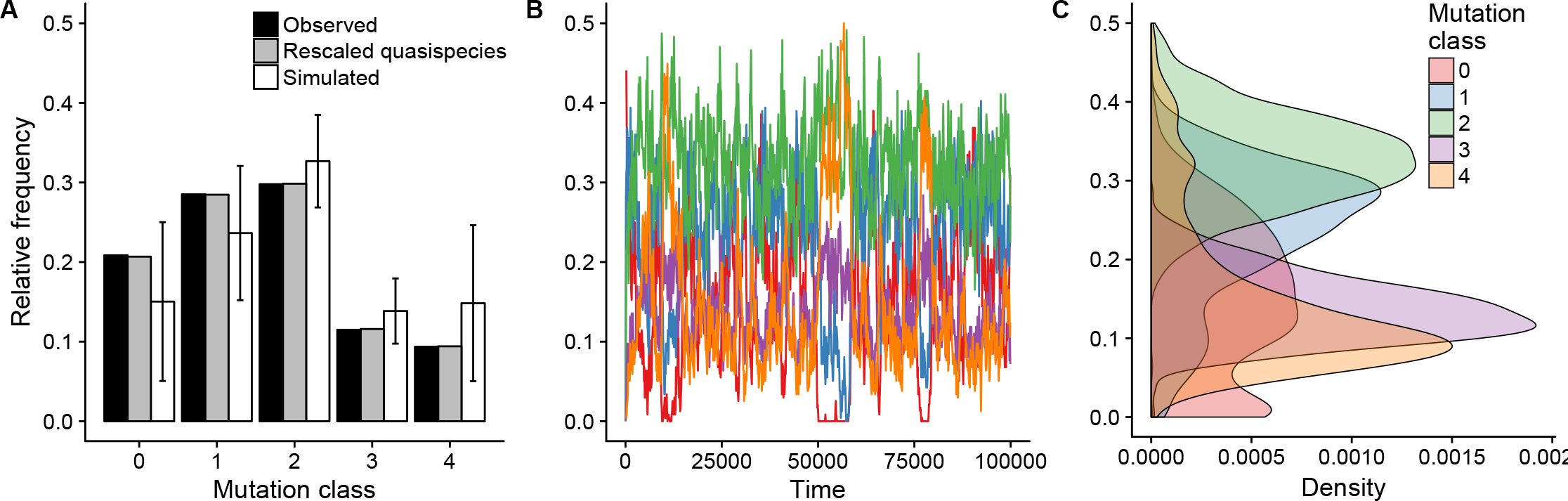
Inferred and observed steady state REPIN frequencies in *P. fluorescens* SBW25. (A) Shows the observed frequencies at a mutation rate of 8.9 × 10^−11^. We rescaled time to allow us to do simulations at a mutation rate of 10^−4^. The resulting quasispecies equilibria agree almost perfectly with the observed frequencies. A simulation of a single Wright-Fisher process (10^5^ generations) with the same fitness values allows us to infer the variation of these frequencies. (B) Relative frequencies obtained from the Wright-Fisher process using the scaled fitness values for 10^5^ generations. (C) Density plot of the relative frequencies of the mutation classes from the Wright-Fisher process.

The rate at which duplications occur can be calculated from the inferred fitness values. We calculate the duplication rate from these fitness values by subtracting one, as “one” is the part of the fitness in our model that corresponds to REPIN maintenance. The duplication rate we inferred for the master sequence in SBW25 is 9.8 × 10^−9^ per generation and per sequence.

However, this means that for the 3^rd^ mutation class, we infer negative duplication rates (Table 1). Unless there is an active deletion process for these mutation classes, these duplication rates are unlikely to be accurate. Alternatively, it is possible that members of the 4^th^ mutation class are more likely to replicate than members of the 3^rd^ mutation class. This could be true as it is possible that these sequences are also recognized by a second RAYT transposase in the SBW25 genome. To alleviate this problem, we can simply scale up all mutation classes so the lowest fitness is 1. This leads to a higher duplication rate of the master sequence’s mutation class of 11.3 × 10^−9^ instead of 9.8 × 10^−9^ (Table 1).

If we assume one cell division to take 40 minutes and novel REPIN insertions to be selectively neutral then it would take about 6734 years until a novel REPIN master sequence fixes in the SBW25 population. This seems to be a surprisingly long time, but it would explain, why, to our knowledge, there is no report of novel REPIN insertions within genomes. It may also explain why REPINs can be maintained for long times within a genome without being selected against because due to the rarity of duplication events the negative fitness effects resulting from transposition (e.g. transposition is likely to disrupt genes because about 88% of the SBW25 genome are coding regions (Silby *et al*. 2009)) are probably negligible.

### REPIN duplication rates in other bacteria

We also calculated duplication rates for four more *Pseudomonas* strains and five more *E. coli* strains. The *E. coli* strains we chose were quite distantly related to each other and belong to phylogroups A, B2 and D. The *Pseudomonas* strains we chose are very distantly related to each other as well as to *E. coli* (Figure 4A). To get an idea about how distantly related the individual strains are, we gauge the time that has passed since the strains diverged by measuring the 16S rDNA divergence (Ochman and Wilson 1987; Ochman *et al*. 1999). Ochman et al. estimated that it takes about 50 million years for the 16S rDNA to diverge by 1%. According to these estimates, the most recent common ancestor (MRCA) of the *E. coli* strains lived approximately 15 million years ago (mya). The MRCA of the *Pseudomonas* strains lived approximately 100 mya and *E. coli* and *Pseudomonas* diverged about 600 mya. Hence, the REPIN populations in our selected bacteria have been evolving independently of each other for a very long time. RAYTs, the genes that mobilize REPINs in *E. coli* and *Pseudomonas*, are also very different in *E. coli* and *Pseudomonas* and belong to two different gene classes (Bertels and Rainey 2011b). There is no detectable sequence conservation in the nucleotide sequence and very little sequence conservation in the aminoacid sequence apart from the catalytic center of the protein.

**Figure 4.**
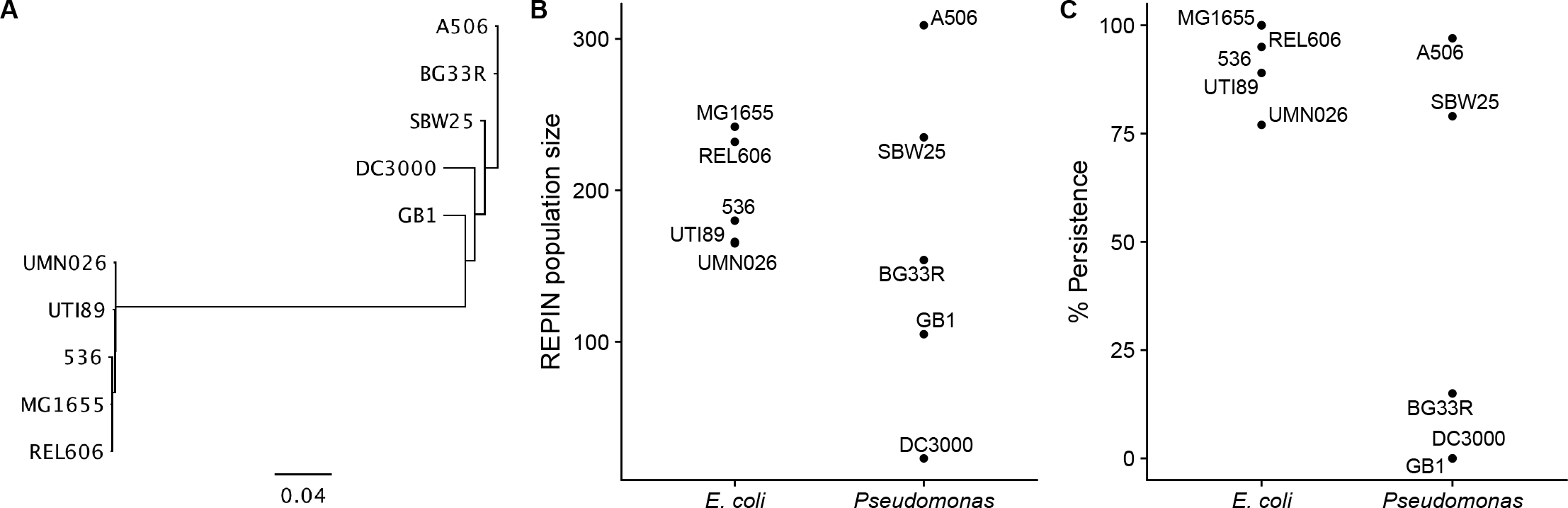
REPIN populations in other bacteria. (A) 16S tree showing the phylogenetic relationship between REPIN containing bacteria selected in our study. The scale bar shows the number of substitutions per nucleotide site. (B) REPIN population sizes in *E. coli* and *Pseudomonas*. (C) Proportion of 100 simulations where at least 10% of all sequences are maintained as master sequence at the end of the simulation.

### Divergent bacteria have divergent REPIN populations

The divergence between the different bacterial strains is also reflected in the similarity between the most abundant 25bp long sequences (REP sequences). The most common sequences in *E. coli* are almost all identical, except for that of UTI89, where the most common sequence is shifted by one nucleotide with respect to the other *E. coli* sequences (File S2). But all *E. coli* REP sequences are very different to all of the *Pseudomonas* REP sequences. Among the *Pseudomonas* strains, the REP sequences from *P. fluorescens* A506 and *P. fluorescens* BG33R are almost identical (again shifted by one nucleotide), which are also the most closely related strains. Despite this similarity, the population sizes and structures are completely different between the two strains (see population networks in File S3). This observation highlights the opportunity to study the evolution of entire populations instead of single strains, which is basically impossible for any other natural population.

REPIN populations in *E. coli* form relatively simple networks, consistent with a single fitness peak. In contrast, REPIN populations from *Pseudomonas* form more complex networks, which is more consistent with a rugged fitness landscape (see sequence networks in File S3). The differences in the complexity of the sequence network may stem from the fact that there is only a single RAYT gene in *E. coli*, but there are usually multiple RAYT genes in *Pseudomonas*. If we assume that the activities of multiple RAYT genes can interfere with each other, then generalist sequences that can be moved by multiple RAYT genes will evolve, and give rise to a complex sequence network.

Although the the divergence between *E. coli* and *Pseudomonas* are very large and the differences between the structure of the REPIN (File S4, whether the REPIN is symmetric as in *Pseudomonas* or not as in *E. coli)* and the corresponding transposase are tremendous (Bertels and Rainey 2011b) the inferred REPIN population sizes are surprisingly similar (Figure 4B). REPIN populations in *E. coli* range between 165 (UMN026) and 242 (MG1655) members. REPIN populations in *Pseudomonas* are spread more widely and range between 23 (DC3000) and 309 (A506) members. The population size has a strong effect on whether the master sequence can persist within the population or whether it will die out. Our simulations show that among all *Pseudomonas* REPIN populations only that of *P. fluorescens* A506 and *P. fluorescens* SBW25 are large enough to persist over long periods of time. In *E. coli*, in contrast, most populations persist over 10^5^ time steps (Figure 4C).

### Small REPIN populations in Pseudomonas

*P. syringae* DC3000 is different from the other *Pseudomonas* strains not only the REPIN population is particularly small (only 23 members), which leads to a particularly unstable REPIN population (Figure 4C). Another notable feature of the DC3000 REPIN population is that a large part of the repetitive sequences does not form REPINs (File S4). This suggests to us that the DC3000 REPIN population may be a dead or dying population, which is slowly disintegrating due to genetic drift. This hypothesis is further supported by the observation that the only RAYT in DC3000 is not flanked by the most common 25bp long sequence in the genome, which is the case for all other population we have analzyed (File S2) and has been a defining feature of the REPIN-RAYT system (Bertels and Rainey 2011b). Together, our data suggests that the reason for the small and unstable REPIN population in DC3000 is that it is slowly disintegrating over time. Hence the population is probably not in equilibrium, which means that the inferred duplication rates may not be accurate.

The populations found in BG33R and GB1 are also too small to persist for extended periods of time. However, in contrast to DC3000, they are also the two populations with the highest inferred duplication rate, and in both cases the most common 25bp long sequence does flank a RAYT gene and both populations consist mostly of REPINs (File S4). Hence there is no sign of population disintegration. The inferred high duplication rates are likely to evolve for small populations, because the mutation load for small populations is comparatively small. This suggests that these two populations may be growing.

### REPIN populations in competition

The population network in BG33R is particularly interesting as it contains two similar sized population (126 and 147 members) and the REPIN master sequence consists in both cases of two identical 25mers that occur both exactly 160 times in the genome and differ in 5 nucleotide positions (i.e. the REPIN master sequence differs in 10 positions). When inferring the fitness of the master sequence for both populations, then we also get very similar and extremely high duplication rates of 32 × 10—9 and 37 × 10^—9^. One would expect the evolution of high duplication rates not only for growing populations but also for populations that are competing for space in the genome. With space we are referring to regions in the genome that are fitness neutral, i.e. regions of the genome that incur no fitness cost when inserted into.

### REPIN populations in E. coli

In *E. coli* the most abundant 25bp long sequences do not form symmetric REPINs as observed in *Pseudomonas* (File S4). This could lead us to the conclusion, as for DC3000, that *E. coli* does not contain any REPIN populations that are alive. However there are a few differences to DC3000. First of all, RAYTs in *E. coli* are very distantly related to RAYTs in most *Pseudomonas*, which leaves the possibility that REPINs in *E. coli* are structured differently to REPINs in *Pseudomonas*. Second, there is not a single instance of a REPIN in any of the five *E. coli* populations. If *E. coli* REPIN populations were dying populations, then all populations in *E. coli* were already dead. This either happened about 15mya, when the last common ancestor of the five *E. coli* strains lived or it happened recently simultaneously. If it happened 15mya, then we would expect the population to have vanished by now and not consist of up to 242 members. It also seems unlikely that it happened recently in all strains at the same time and within the same time frame all the REPINs vanished but the singlets remained. Finally, the most common 25bp long sequences in the five strains does still flank the RAYT gene something that is not the case for DC3000 but for all other REPIN populations in our study (File S2).

### REPIN duplication rate is close to the error threshold

The duplication rates of the master sequences are in the range of 5 × 10^−9^ and 37 × 10^−9^ and 5 × 10^−9^ and 15 × 10^−9^ when excluding unstable populations. Considering that the rates were inferred for very different species and the species contain very different transposases that disperse the REPIN populations, these values are very similar. This may be due to at least two reasons. First, the duplication rate is very close to its lower possible limit, because the number of mutations that occur on average between two duplication events is between 0.12 and 0.39 for *Pseudomonas* (0.29 and 0.39 without unstable populations) and between 0.22 and 0.46 for *E. coli* (0.39 and 0.46 without UMN026). If on average one mutation occurs between two duplication events, then it is impossible to maintain a master sequence. For our model a master sequence cannot be maintained above a frequency of 1% when the duplication rate of the master sequence and all other sequences is equal or lower than 2.2 × 10^−9^ for *E. coli* and 4.4 × 10^−9^ for *Pseudomonas* (File S5 and Figure 5). Second, each duplication event can be seen as a mutation that is introduced at a random position in the genome. This means that an increase in the duplication rate would also increase the mutational load for the host organism. Hence, similar to selection for replication fidelity (Lynch *et al*. 2016), selection will favor organisms with decreased REPIN duplication rates, but is limited by the power of random genetic drift. The REPIN duplication rates we inferred are probably the result of these two opposing forces.

**Figure 5.**
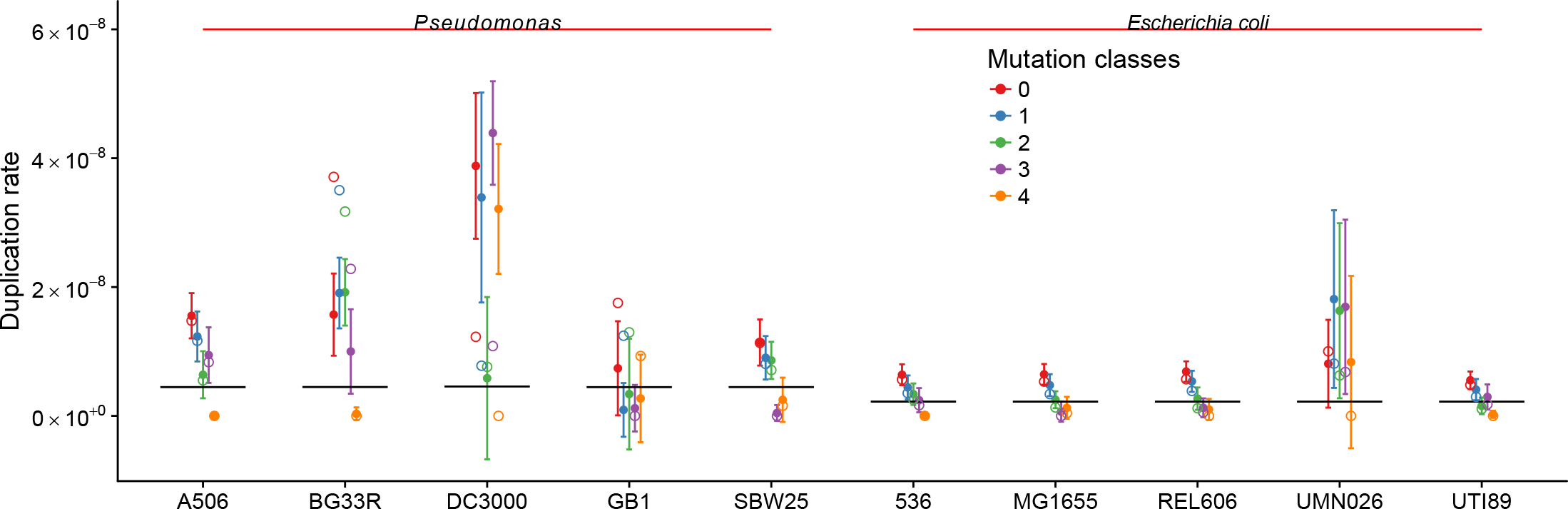
REPIN duplication rates in *Pseudomonas* and *E. coli* strains. The figure shows duplication rates for the largest REPIN populations in various *Pseudomonas* and *E. coli* strains. The solid circles indicate the mean duplication rate and their variance inferred from the frequencies of the aforementioned Wright-Fisher-Process at 20 random positions of the simulation. For BG33R, DC3000, GB1 and UMN026 values from the simulation are not reliable as the master sequence did not persist until the end of the simulation. Empty circles indicate the inferred duplication rate from the observed sequences. The black lines indicate error thresholds. If the duplication rate of the master sequence falls below the black horizontal lines, then it is impossible to maintain the master sequence above a frequency of 1% in the population. All error thresholds among *Pseudomonas* strains and among *E. coli* strains only differ at a level of 10^−10^, which cannot be seen in the figure as it is less than the line width. The full organism names from left to right are: *P. fluorescens* A506, *P. synxantha* BG33R, *P. syringae* pv. *tomato* DC3000, *P. fluorescens* SBW25, *P. putida* GB1, *E. coli* 536, *E. coli* K-12 MG1655, *E. coli* B REL606, *E. coli* UMN026, *E. coli* UTI89.

### Maintenance of the REPIN-RAYT system

The low duplication rate we inferred for all REPIN populations also suggests that REPIN sequences have been part of bacterial genomes for a very long time. This again raises the question of how and why they are maintained. There are two explanations: (1) the REPIN-RAYT system is frequently transmitted horizontally or (2) they provide a benefit to the host organism (Bichsel *et al*. 2013).

It is possible that the REPIN-RAYT system does get horizontally transferred from time to time. However, horizontal transfers are likely to be rare, because in order to establish a novel REPIN population in a new host both the transposase (RAYT) and the REPIN have to be transferred. This process is probably facilitated by the fact that RAYTs are usually flanked by REPINs (Bertels and Rainey 2011b). However, the rarity of these events is consistent with the observation that the establishment of a population that is as diverse as the REPIN population in SBW25 will take thousands of years. Hence it seems unlikely that horizontal transfers are frequent enough to explain the ubiquitous presence of the REPIN-RAYT system in bacteria.

Alternatively, the REPIN-RAYT system may be maintained because it provides a selective advantage to the host bacterium. For individual REP sequences there have been many studies on potential benefits (Liang *et al*. 2015; Higgins *et al*. 1988; Espéli *et al*. 2001). However, local benefits are unlikely to outweigh the detrimental effects of transposition into genes or regulatory regions let alone explain the maintenance of the REPIN-RAYT system. It seems more likely that the REPIN-RAYT system possesses a function other than the dispersion of REPINs that is beneficial for the host bacterium.

## Acknowledgements

All authors acknowledge the generous funding from the Max Planck Society.

